# High-resolution app data reveal sustained increases in recreational fishing effort in Europe during and after COVID-19 lockdowns

**DOI:** 10.1101/2022.12.07.519488

**Authors:** Asta Audzijonyte, Fernando Mateos-González, Justas Dainys, Casper Gundelund, Christian Skov, J. Tyrell DeWeber, Paul Venturelli, Vincentas Vienožinskis, Carl Smith

## Abstract

It is manifest that COVID-19 lockdowns extensively impacted human interactions with natural ecosystems. One example is recreational fishing, an activity which involves nearly 1 in 10 people in developed countries. Fishing licence sales and direct observations at popular angling locations suggest that recreational fishing effort increased substantially during lockdowns. However, the extent and duration of this increase remain largely unknown due to a lack of objective data. We used four years (2018 to 2021) of anonymous, high-resolution data from a personal fish-finder device to explore the impact of COVID-19 lockdowns on recreational fishing effort in four European countries (Lithuania, the Czech Republic, Denmark, and Germany). We show that device use and, by extension, angling effort increased 1.2-3.8 fold during March-May 2020 and remained elevated even at the end of 2021 in all countries except Denmark. Fishing during the first lockdown also became more frequent during weekdays. Statistical models with the full set of fixed (weekdays, lockdown, population) and random (season, year, administrative unit) factors typically explained 50-70% of the variation, suggesting that device use and angling effort were relatively consistent and predictable through space and time. Our study demonstrates that recreational fishing behaviour can change substantially and rapidly in response to societal shifts, with profound ecological, human well-being and economic implications. We also show the potential of angler devices and smartphone applications to supply data for high-resolution fishing effort analysis and encourage more extensive science and industry collaborations to take advantage of this information.

**Significance statement:** Recreational fishing is a popular and widespread activity with ecological, social and economic impacts, though problematic to assess and manage due to a paucity of information regarding effort and catch. Here, we use high-resolution data from a personal angler sonar device to show how the COVID-19 pandemic changed angler behaviour and fishing effort across Europe. We demonstrate that angling effort doubled and remained higher at the end of 2021 than before the first lockdowns. Such rapid and profound changes could have significant consequences for aquatic ecosystems, possibly requiring new management approaches. We encourage the adoption of novel data from angler devices, citizen science, and more active science-industry collaborations to improve recreational fishing assessment and management.

## Introduction

The COVID-19 pandemic and associated lockdowns affected the natural world and provided novel research opportunities. In many cases, the slowing of human activities, coined the “Anthropause” (1), had positive effects on nature through reduced traffic (2, 3), noise and other pollution (4, 5), airspace fragmentation and human activity in coastal areas (6–8) (but see Bates et al. (9) for the full range of impacts). There is also evidence that the Anthropause had a positive effect on marine fishes (10, 11) through reductions in small-scale (12–15) and, to a lesser degree, large-scale commercial fisheries (16). Commercial seafood catches in the US alone were estimated to be 40% lower in 2020 compared to 2019 (17).

Although the positive effects of a reduction in commercial fishing are reasonably clear, impacts from recreational fishing are more opaque (9, 18). This uncertainty stems from a lack of temporally and spatially resolved data for recreational fishing effort. Even in cases where these data are available, they are often based on fishing licence sales, creel surveys, questionnaires or expert knowledge that may not reflect changes in angling behaviour, especially in atypical circumstances. Nevertheless, existing lines of evidence provide intriguing insights into recreational fishing behaviour during government imposed lockdowns. For example, a survey of diary panellists in the UK suggested that the fishing effort of sea anglers declined during 2020 in response to the lockdown (19). In contrast, the sale of angling licences in Denmark increased by 20% in March-May of 2020 (20), first-time or returning anglers increased by 20% in Ontario, Canada (21), fishing trips increased 20% in the United States (21), and apparent illegal fishing increased ∼10-fold in some marine-protected areas in British Columbia, Canada (22). A more nuanced pattern was observed in Western Australia where metropolitan boat anglers seemingly fished less due to travel restrictions, but anglers outside of urban centres increasing the frequency of fishing trips (23). Overall, these studies suggest that that changes in recreational fishing effort and participation might have been country-specific because different countries imposed different types of restrictions. For example, access to fishing was restricted for the British, whereas Danes were encouraged to spend time outdoors.

Recreational fishing constitutes the major or even sole source of fishing mortality in many freshwater and coastal fish stocks (24–26). In addition, angling can contribute substantially to local economies (27), represents an important source of protein (28) and provides important well-being and health benefits (29). Recreational fisheries are potentially more sensitive to social and societal changes than commercial fisheries, and major changes in recreational fishing are likely to have important consequences for target fish species, local economies, and angler health. The COVID-19 pandemic provides an important opportunity to understand angler behaviour - not only in response to rare events like an Anthropause, but also as a natural experiment that can provide insights into more general aspects of human behaviour in the context of recreational fisheries management. In this study, we used anonymous personal sonar device data as a proxy for recreational angler fishing effort in four European countries with different socioeconomic and urbanisation levels and different recreational fishing management rules (Lithuania, Czech Republic, Denmark, and Germany). We asked three main questions. First, did angling effort change during Covid-19 lockdowns, and were there common patterns among countries? Second, if changes occurred, were they sustained once lockdowns ended? Third, which factors explain the estimated angling effort through space and time, and can our statistical models explain this effort distribution?

## Results

Recreational fishing effort in the Czech Republic, Denmark, Germany, and Lithuania was assessed using anonymous data harvested from a widely used low cost portable fish finder device *Deeper Sonar* (https://deepersonar.com/). Recreational anglers use this device to locate fish, measure water depth, and create small-scale bathymetric maps as an aid to angling. A recent study calibrated the proportion of anglers that used this sonar device at a popular fishing location in Lithuania during 2020-2021 (30), and showed high correspondence between the proportion of device users and the actual number of anglers on a given day (95% posterior probabilities of 1.5-2.6% of anglers). These findings imply that angler behaviour and total recreational fishing effort may be reasonably well predicted from sonar usage, although the quality of this prediction is likely to vary among countries (see Discussion). We explored daily angler sonar usage counts per municipality, county, or other administrative unit, as appropriate, in each of the four countries, and assessed how this usage compared before, during, and after COVID-19 lockdowns, using the lockdown stages and definitions provided by Our World in Data compilation (Fig. S1). Depending on the country, the number of unique sonar users by the end of 2021 constituted 0.35-10% of the estimated total anglers in the country (Table 1). We assumed that changes in sonar device use reflected changes in the overall angling effort, but see Discussion for further clarifications and caveats.

**Table 1.** Summary data for each of the four countries included in the study, with lockdown phases taken from Our World in Data compilation of stay-at-home requirement in the ‘Policy Responses to the Coronavirus Pandemic’, and government data for Denmark (see Supplementary material). The total number of unique sonar users and fishing trips recorded is shown for the last day of the phase. The 2020 population of each country is shown in millions (71). For details on angler number estimation see Methods. Percentage shows the proportion of unique sonar device users at the end of the study period relative to the total estimated number of anglers.

| Country<br>(population<br>size in 2020) | Lockdown phase | Start | End | Days | Unique<br>users | Trips<br>recorded | Estimated<br>angler<br>numbers |
| --- | --- | --- | --- | --- | --- | --- | --- |
| **Czech**<br>**Republic**<br>**(10.7M)** | Pre lockdown | 2018-01-01 | 2020-03-14 | 804 | 969 | 15437 | 320 K |
|  | Lockdown 1 | 2020-03-15 | 2020-05-31 | 77 | 3957 | 19232 | (~2%) |
|  | Post lockdown 1 | 2020-06-01 | 2020-10-21 | 142 | 4773 | 26210 |  |
|  | Lockdown 2 | 2020-10-22 | 2021-04-11 | 171 | 5382 | 29843 |  |
|  | Post lockdown 2 | 2021-04-12 | 2021-12-31 | 264 | 6979 | 42297 |  |
| **Denmark**<br>**(5.8M)** | Pre lockdown | 2018-01-01 | 2020-03-10 | 800 | 830 | 2909 | 442 K |
|  | Lockdown 1 | 2020-03-11 | 2020-05-31 | 81 | 891 | 3317 | (~0.35%) |
|  | Post lockdown 1 | 2020-06-01 | 2020-10-22 | 143 | 1153 | 4662 |  |
|  | Lockdown 2 | 2020-10-23 | 2021-05-20 | 209 | 1327 | 5643 |  |
|  | Post lockdown 2 | 2021-05-21 | 2021-12-31 | 225 | 1568 | 7035 |  |
| **Germany**<br>**(83.8M)** | Pre lockdown | 2018-01-01 | 2020-03-08 | 798 | 10866 | 29294 | 6 570 K |
|  | Lockdown 1 | 2020-03-09 | 2020-05-05 | 57 | 12349 | 37405 | (~0.38%) |
|  | Post lockdown 1 | 2020-05-06 | 2020-10-14 | 161 | 17159 | 68498 |  |
|  | Lockdown 2 | 2020-10-15 | 2021-10-09 | 359 | 24410 | 118551 |  |
|  | Post lockdown 2 | 2021-10-10 | 2021-12-31 | 83 | 25363 | 125955 |  |
| **Lithuania**<br>**(2.7M)** | Pre lockdown | 2018-01-01 | 2020-03-15 | 805 | 12573 | 68505 | 250 K |
|  | Lockdown 1 | 2020-03-16 | 2020-06-16 | 92 | 14032 | 81381 | (~10%) |
|  | Post lockdown 1 | 2020-06-17 | 2020-08-06 | 50 | 14751 | 89338 |  |
|  | Lockdown 2 | 2020-08-07 | 2021-04-18 | 254 | 18140 | 129740 |  |
|  | Post lockdown 2 | 2021-04-19 | 2021-12-31 | 257 | 20348 | 157254 |  |

Generalised linear mixed model showed that COVID-19 lockdowns had substantial and relatively consistent effects on estimated recreational angling effort in three of four study countries. The first lockdown phase in boreal spring of 2020 lasted between 57 (Germany) to 81 (Denmark) days, and was generally associated with substantial and significant increases in the estimated effort. Effort in the Czech Republic, Germany and Lithuania increased 2.2-3.8 times (P <0.001, Fig. 1, Tables 2,3), and remained substantially higher (1.7-3.4 times higher than before the pandemic) even after the first lockdown in summer and autumn of 2020. The second lockdown phase broadly corresponded with the boreal autumn and winter of 2020-2021, when fishing effort tends to be low relative to spring and summer (see Fig. S2-5 for the plots on random model effects). This seasonal variation in effort was accommodated by random effects in the model, and the results show that fishing effort during the second lockdown in Czech Republic, Germany and Lithuania was still much higher than before the pandemic. Germany was an exception in that the second lockdown phase lasted almost a year, but fishing effort during this period remained 1.3-2.5 times higher than before the pandemic. The increased fishing effort was sustained in all three counties even after all lockdowns ended in mid-2021 (1.2-2.8 times higher than before the pandemic). Denmark showed a slightly different pattern to the other three countries. There was about a 20% increase in effort during the March-May of 2020 (P = 0.023), with a return to pre-pandemic effort levels during the summer of 2020 (first post lockdown period) and strong indications of a 50-60% reduction in effort relative during the second lockdown and second post-lockdown phase in comparison with the pre-lockdown stage.

**Table 2.** (can be moved to the supplementary material). Summary of fixed effects for negative binomial GLMMs fitted to sonar user data for the four countries investigated in the study. Results show the impact of phase of COVID-19 lockdown (Pre lockdown, During lockdown 1, Post lockdown 1, During lockdown 2, Post lockdown 2), day of week (Weekday, Weekend), regional population size, number of unique sonar users and interactions between phase of lockdown and day of week. The pre lockdown phase and weekday are the intercept of the models. The marginal and conditional R^2^ indicate the proportion of variance in sonar usage data explained by the model fixed effects and fixed + random effects, respectively. Details on model random effects are shown in Figs S3-6 and Table S1.

| Coefficient | Country |  |  |  |  |  |  |  |  |  |  |  |
| --- | --- | --- | --- | --- | --- | --- | --- | --- | --- | --- | --- | --- |
|  | Czech Republic (N = 20,454) |  |  | Denmark (N = 140,256) |  |  | Germany (N = 546,414) |  |  | Lithuania (N = 78,184) |  |  |
|  | *Estimate* | *95% CI* | *P-value* | *Estimate* | *95% CI* | *P-value* | *Estimate* | *95% CI* | *P-value* | *Estimate* | *95% CI* | *P-value* |
| Intercept | -0.11 | -0.95 – 0.72 | 0.793 | -3.65 | -4.04 – -3.01 | <0.001 | -3.07 | -3.51 – -2.62 | <0.001 | -0.83 | -1.14 – -0.36 | <0.001 |
| Day(Weekend) | 0.50 | 0.46 – 0.54 | <0.001 | 0.68 | 0.60 – 0.76 | <0.001 | 0.59 | 0.56 – 0.62 | <0.001 | 0.97 | 0.95 – 0.99 | <0.001 |
| Population size | 0.35 | -0.02 – 0.72 | 0.061 | 0.38 | 0.18 – 0.59 | <0.001 | -0.18 | -0.33 – -0.02 | 0.024 | 0.10 | -0.12 – 0.33 | 0.379 |
| Unique users | 0.57 | 0.46 – 0.67 | <0.001 | 0.46 | 0.24 – 0.69 | <0.001 | 0.46 | 0.41 – 0.52 | <0.001 | -0.19 | -0.22 – -0.10 | <0.001 |
| Phase(Lockdown1) | 0.80 | 0.68 – 0.91 | <0.001 | 0.20 | 0.03 – 0.38 | 0.023 | 1.00 | 0.93 – 1.06 | <0.001 | 1.33 | 1.27 – 1.39 | <0.001 |
| Phase(Post1) | 0.53 | 0.42 – 0.64 | <0.001 | -0.08 | -0.28 – 0.12 | 0.434 | 1.11 | 1.05 – 1.17 | <0.001 | 1.23 | 1.16 – 1.30 | <0.001 |
| Phase(Lockdown2) | 0.28 | 0.15 – 0.40 | <0.001 | -0.91 | -1.17 – -0.65 | <0.001 | 0.69 | 0.62 – 0.76 | <0.001 | 0.92 | 0.85 – 0.98 | <0.001 |
| Phase(Post2) | 0.72 | 0.56 – 0.88 | <0.001 | -0.83 | -1.16 – -0.50 | <0.001 | 0.16 | 0.08 – 0.25 | <0.001 | 1.03 | 0.95 – 1.12 | <0.001 |
| Phase(Lockdown1) x Day(Weekend) | -0.17 | -0.26 – -0.07 | <0.001 | -0.16 | -0.32 – 0.01 | 0.065 | -0.13 | -0.18 – -0.07 | <0.001 | -0.36 | -0.41 – -0.30 | <0.001 |
| Phase(Post1) x Day(Weekend) | -0.12 | -0.19 – -0.04 | 0.002 | -0.14 | -0.28 – -0.01 | 0.040 | -0.22 | -0.25 – -0.18 | <0.001 | -0.32 | -0.39 – -0.26 | <0.001 |
| Phase(Lockdown2) x Day(Weekend) | 0.26 | 0.17 – 0.35 | <0.001 | 0.26 | 0.10 – 0.41 | 0.001 | 0.00 | -0.03 – 0.03 | 0.943 | -0.04 | -0.08 – 0.01 | 0.099 |
| Phase(Post2) x Day(Weekend) | -0.04 | -0.10 – 0.02 | 0.196 | -0.25 | -0.39 – -0.11 | <0.001 | 0.12 | 0.07 – 0.18 | <0.001 | -0.19 | -0.24 – -0.16 | <0.001 |
| Marginal/Conditional $R^2$ | 0.30 / 0.70 | | | 0.06 / 0.30 | | | 0.11 / 0.51 | | | 0.14 / 0.53 | | |

**Table 3.** Total number of predicted sonar user trips for the four countries for each of the lockdown periods based on negative binomial GLMMs. Because the number of trips depend on the population size and the number of unique users, these predictions use average population sizes and average number of unique users in each country. Results show predicted total number of user trips for each of the studied time periods with and without the lockdown effect included, and their ratio. Numbers in parenthesis show 95% confidence intervals. For all calculation details see Excel spreadsheet in https://github.com/astaaudzi/covid_angling

| Country | Lockdown phase | Number of trips | Number of trips without lockdowns | Ratio |
| --- | --- | --- | --- | --- |
| Czech Republic | Pre lockdown | 11,947 (5077-27841) | - | - |
|  | Lockdown 1 | 2,389 (875-6510) | 1,074 (433-2594) | 2.2 (2.0-2.5) |
|  | Post lockdown 1 | 3,410 (1270-9126) | 2,007 (834-4812) | 1.7 (1.5-1.9) |
|  | Lockdown 2 | 3,763 (1346-10355) | 2,844 (1159-6941) | 1.3 (1.2-1.5) |
|  | Post lockdown 2 | 7,916 (2806-22158) | 3,853 (1603-9191) | 2.1 (1.8-2.4) |
| Denmark | Pre lockdown | 2904 (1668-4991) | - | - |
|  | Lockdown 1 | 336 (154-742) | 275 (149-507) | 1.2 (1.0-1.5) |
|  | Post lockdown 1 | 448 (201-991) | 486 (266-879) | 0.9 (0.8-1.1) |
|  | Lockdown 2 | 344 (141-835) | 855 (454-1600) | 0.4 (0.3-0.5) |
|  | Post lockdown 2 | 321 (127-810) | 737 (405-1336) | 0.4 (0.3-0.6) |
| Germany | Pre lockdown | 17,037 (10837-27060) | - | - |
|  | Lockdown 1 | 3,103 (1812-5344) | 1,142 (715-1851) | 2.7 (2.5-2.9) |
|  | Post lockdown 1 | 9,543 (5667-16306) | 3,145 (1983-5061) | 3.0 (2.9-3.2) |
|  | Lockdown 2 | 15,265 (8944-26339) | 7,657 (4811-12318) | 2.0 (1.9-2.1) |
|  | Post lockdown 2 | 2,176 (1250-3887) | 1,854 (1154-3027) | 1.2 (1.1-1.3) |
| Lithuania | Pre lockdown | 29,887 (21698-48315) | - | - |
|  | Lockdown 1 | 10,807 (7259-18983) | 2,858 (2039-4728) | 3.8 (3.6-4.0) |
|  | Post lockdown 1 | 5,356 (3536-9499) | 1,566 (1108-2589) | 3.4 (3.2-3.7) |
|  | Lockdown 2 | 23,228 (15417-40910) | 9,257 (6590-15354) | 2.5 (2.3-2.7) |
|  | Post lockdown 2 | 24,232 (15897-43422) | 8,651 (6148-14168) | 2.8 (2.6-3.1) |

**Figure 1.**
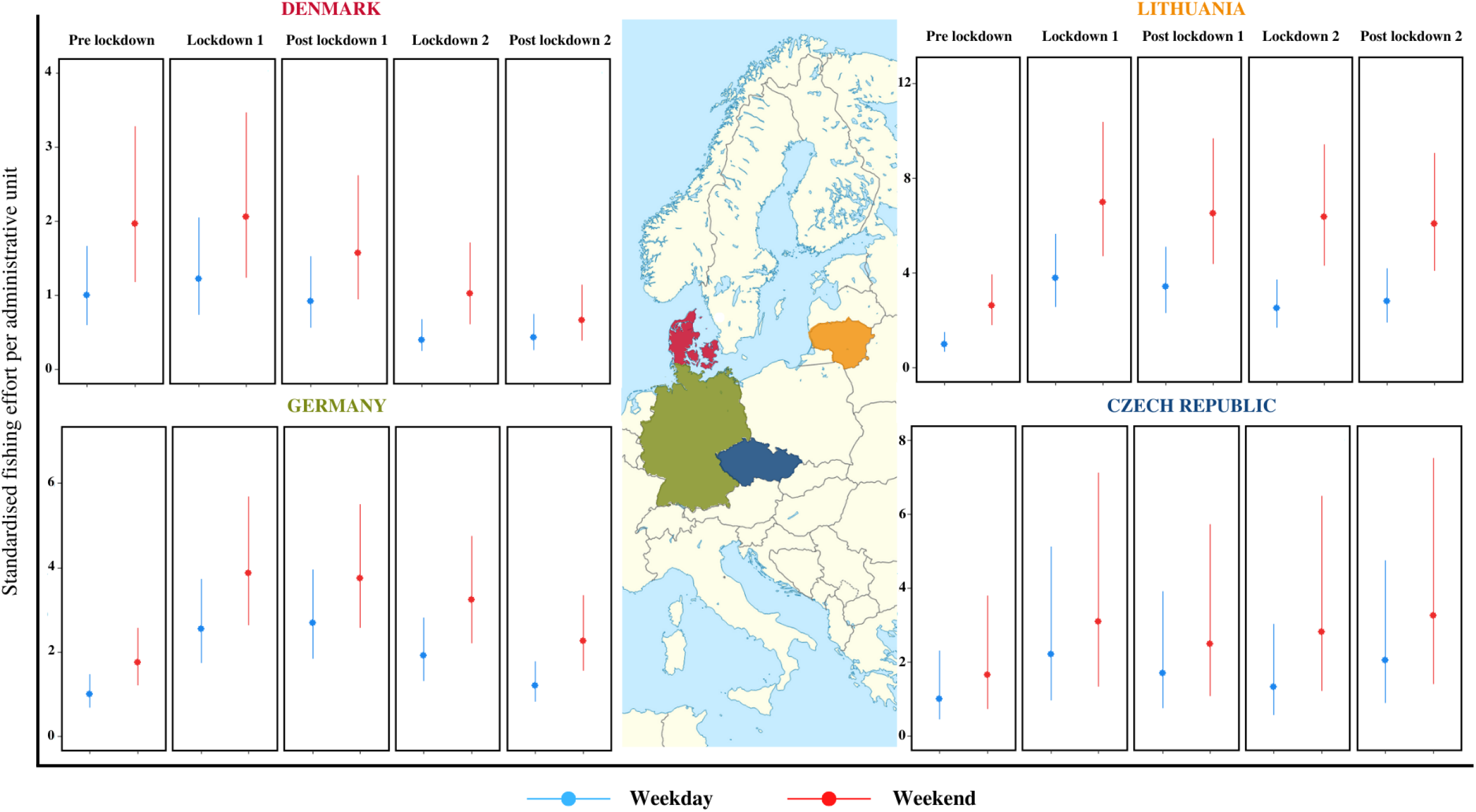
Estimated effort per administrative unit (as number of sonar buoy users during weekdays and weekends) by country for each study period (Pre lockdown, During lockdown 1, Post lockdown 1, During lockdown 2, Post lockdown 2). Effort was estimated for an average administrative unit size for each country, scaled to the weekday of the Pre lockdown stage. For effort predictions across all population sizes see Fig. S8.

As expected, recreational fishing effort was considerably higher during weekends compared to weekdays. This difference likely reflected both the real difference in fishing effort, and a higher probability of device use during weekends, since the device may be more popular among employed anglers with higher incomes (26). Notably, one of the most consistent features of fishing activity during the first lockdown stage was the significant negative interaction between effort and weekend (Table 2), indicating a more even distribution of fishing activities during the week (Fig. S6, Table 2). The pattern was less consistent during the second lockdown, where the difference in effort distribution increased compared to the pre-pandemic levels in the Czech Republic and Denmark (P <0.001), but not in Germany or Lithuania (Table 2).

In addition to assessing changes in recreational fishing effort during the COVID-19 pandemic, our models explained a surprisingly large proportion of variance (51-70%) through space and time in Germany, Czech Republic, and Lithuania (conditional R^2^ in Table 2). This result suggests that the temporal and spatial distribution of device use and, by inference, recreational fishing effort, was consistent and predictable. The models explained less variation for Denmark (31%), possibly due to the low uptake of the device in this country (estimated 0.35% of total anglers, Table 1). The fixed effects of lockdown phase, weekend and population density explained 11-30% of the variation in Germany, Czech Republic and Lithuania, and 6% in Denmark. The remaining variance (40% in Czech Republic, 24% in Denmark, 40% in Germany and 39% in Lithuania) was explained by the random effects, comprising administrative unit, year and season (Fig. S2-5, Table 2). With the exception of Germany, predicted angling effort tended to be greater in municipalities or counties that had larger population sizes, although the effect was only significant in Denmark. This observation suggests that anglers were more likely to fish within a short distance of their homes, rather than travel in search of fishing opportunities. In contrast, the significant negative association between population size and angling effort in Germany (P = 0.024, Table 2) indicates that more angling occurs in less populated areas.

## Discussion

Our study demonstrated substantial and largely consistent COVID-19 lockdown-driven changes in recreational fishing effort across four European countries with different socioeconomic conditions and fishing regulations. The estimated fishing effort in Lithuania, Germany, and Czech Republic increased 2- to 3-fold during the first lockdown of spring 2020 (first research question) and became more evenly distributed during the week. Notably, increases in recreational fishing effort persisted to the end of 2021, almost two years after the imposition of the first lockdown (second research question). By the end of 2021, this increase remained about two times higher than the pre-pandemic period in Lithuania and Czech Republic, and ∼10-30% higher in Germany. The persistence of elevated fishing effort hints at extensive and long term societal changes resulting from COVID-19 lockdowns. Such changes are widely documented in economic systems (31), healthcare (32) and travel (33), but are less recognised in the context of outdoor recreation and potential human impacts on the natural world. Anecdotal evidence and published studies have already suggested that the COVID-19 pandemic drove large increases in recreational fishing activity, especially in the developed world (9, 20–22, 34). However, none of these studies had sufficiently long time series of data to assess whether changes were sustained in the medium term. To our knowledge, ours is the first study to document recreational effort changes using daily data from across four countries for an extended period.

The 2- to 3-fold increase in recreational fishing effort that we observed was considerably higher than estimates from other studies, e.g. 20% increases in Denmark (20), Canada (34) and the United States (21). Moreover, while responses across smaller spatial and temporal scales may be complex (e.g., Ryan et al. (23) in Western Australia), we show that when analysed on a larger scale, human behaviour changes were surprisingly consistent across countries, despite different COVID-19 regulations and recreational fishing traditions. This consistency suggests that similar changes in recreational fishing have likely occurred in other developed countries, and perhaps globally. The most consistent response occurred during the first lockdown in March-May of 2020, with an increase in effort across all four studied countries and relatively more fishing occurring during weekdays. The first lockdown was the period with the most consistent and stringent restrictions and policies. Many businesses and schools closed, and lockdown measures were strictly enforced. Policies during the subsequent phases of post - lockdown and second lockdown were move variable, which likely translated into greater among-country variation in the intensity and weekly distribution of fishing activity. Interestingly, observed recreational effort appeared to track EU unemployment data, suggesting a potential secondary effect of COVID-19 on recreation through job losses and increased free time (35) (see Fig S7).

Our estimates of recreational fishing effort in Denmark were unique in that we observed a relatively small, ∼20% initial increase (95% CI range of 0-50%, Table 3) in effort, followed by a subsequent decrease in 2021 to levels lower than before the pandemic. The initial increase during the spring 2020 lockdown was similar in magnitude to the increase in fishing licence sales reported in Gundelund and Skov (20). The more relative increase in weekday and decrease in weekend angling effort was also consistent to what has been found among users of the Fangstjournalen digital citizen science platform (20). It is harder to explain the apparent large decease in the estimated fishing effort in the post - lockdown period and during 2021. Interestingly, the sale of the mandatory annual angling licenses in Denmark was indeed lower in 2021 (151,686 licenses) compared to 2020 (169,931 licenses), but still well above the situation immediately pre-COVID; e.g., in 2018 (135,082 licenses) and 2019 (137,133 licenses) (36). Hence, both the sonar device and license sales data support the observation that an increase of recreational angling in 2020 was not sustained in 2021, but they paint a different picture about the decrease in effort during 2021 compared to the pre-pandemic levels. On one hand, this discrepancy could be driven by a low uptake of sonar device in Denmark. The absolute number of device users in Denmark is considerably lower than in other countries (Table 1), which could make the data more influenced by the behaviour of a subset of active users, especially if they are responsible for a significant proportion of total trips. On the other hand, it is possible that the anglers who bought annual angling licenses in 2021 intended to fish more than during the pre-covid periods, but ultimately went on fewer fishing trips. Resolving this discrepancy requires data for calibrating sonar device usage to the actual number of anglers in Denmark, as was done in Lithuania (30) and for the citizen science platform Fangstjournalen users in Denmark (37).

Given the difficulties of estimating recreational fishing effort directly, many studies use fishing licence sales as an indication of effort or even the only source of data (21, 37–39). Yet, the discrepancy that we observed between annual licence sales and the number of fishing trips estimated with the sonar device usage is not unique to Denmark. For example, the large increase in effort observed in Lithuania was not reflected in the annual angling sales during 2020-2021 (76k on average) compared to the previous periods (74k on average, during 2015-2018), and only small changes in monthly (19k vs. 21k; same periods) or 2-day licence (14k vs. 16k) sales. Licence numbers are even harder to track in Germany, where licences are sold for different water bodies separately by various organisations and fishing clubs, and there is no national register. These observations indicate considerable difficulties in obtaining accurate estimates of recreational angling effort, given that anglers in most countries are not required to report individual fishing trips. The challenge is even greater during periods of societal shifts when human behaviour is changing quickly, and existing data collection methods are not equipped to track these changes. Our study demonstrates an alternative approach to estimating recreational effort in near real time and at high temporal and spatial resolutions. However, our approach assumes that sonar usage reflects recreational fishing effort, which may not be the case if sonar users behave differently from the majority of anglers, or if the usage of the device itself changed during the pandemic (e.g., anglers were more likely to use it). Data could also be biased if device use is more popular in certain types of fishing than others, which is likely to be the case. Yet, the overall consistency of our findings with general expectations of effort change through different lockdown phases (initial increase and subsequent decrease) suggests that the biases might be reasonably small, and the device usage may represent a reliable proxy of the total effort.

Our study is not the first to use novel digital data to assess human activities and recreational fishing. Smartphone applications and other digital methods, e.g., web scraping, provide an increasingly common approach to explore fishing effort and catches. These methods are different to the traditional data collection techniques (e.g., creel surveys, recall surveys, boat ramp and aerial surveys), which aim for random sampling and probabilistic methods (40). In contrast, app and other digital methods are often based around self-selection, which makes them non-probabilistic and likely biased (41). Although probabilistic surveys generally outperform non-probabilistic methods, they can be complex and expensive to conduct and are still susceptible to bias (42–45). Moreover, digital approaches to angler effort have often provided effort estimates that are similar to those of traditional survey methods. For example, Papenfuss et al. (46) found a positive relationship between estimated (creel) and observed (app) trips in summer in Alberta, Canada. Similarly, Gundelund et al. (37) found a significant relationship between anglers counted during aerial surveys and active users of the Fangstjournalen platform, which indicated that ∼10 % of the observed anglers were citizen science participants. Data from the electronic citizen science platform MyCatch provided similar patterns of regional fishing activity and spatial distribution of participants compared to traditional survey methods in Alberta, Canada (47). Further, 40 aerial surveys of the Kaunas Reservoir in Lithuania in 2020 showed that the number of the Deeper sonar users, the device used in this study, accurately reflected total angler numbers with narrow Bayesian posterior probability ranges (95% posterior probability of 1.5-2.6% of anglers using the device during the open water season) (30). These studies point to the potential of electronic angler platforms for providing high-resolution spatial and temporal data in near real time, which is important and timely, given the growing popularity of recreational fishing and paucity of data for estimating recreational effort and catches in most countries (48). However, to ensure that electronic data platforms provide reliable and widely acceptable results, we urgently need validation and calibration studies to estimate possible biases and monitor changes in device usage through time.

To address our third question, regarding factors that explain the estimated angling effort through space and time, we note that statistical models applied here explained a surprisingly high proportion of variance, given the inherently noisy nature of human behavioural data (Table S1, Fig. S2-5). We also observed a positive association between estimated angling effort and the human population density in Denmark, Lithuania, and Czech Republic, contrasting with a negative association in Germany. This association is evident in Figs. 2 and S8, although it was not identified as significant in our models for Lithuania and Czech Republic, because of a high correlation with the municipality random effect (see Methods, Table 2). A positive association with population size might indicate that many anglers fish close to their homes and that fish populations in densely populated areas are likely to be under higher recreational fishing pressure compared to more remote locations. While this is expected in large countries like Canada and the USA (although see Weir et al. (49) for evidence of high angler movement rates in USA), anglers in smaller countries may be more likely to travel large distances to visit the most popular fishing locations. The negative association between population size and angling effort in Germany was unexpected. We believe that it could have been driven by greater industrialisation in Germany, where densely populated areas may have fewer attractive fishing opportunities. Given that the administrative units in Germany are relatively small, it is likely that many urban anglers travel to neighbouring, less developed, areas for angling (50). This relocation of angling effort is easier in parts of the country with large angler associations, as agreements often allow anglers from nearby states to purchase discounted permits for association waters. The decreasing acceptance of angling as a recreational activity (51) may also drive anglers to rural areas where potential confrontations may be less likely.

**Fig. 2.**
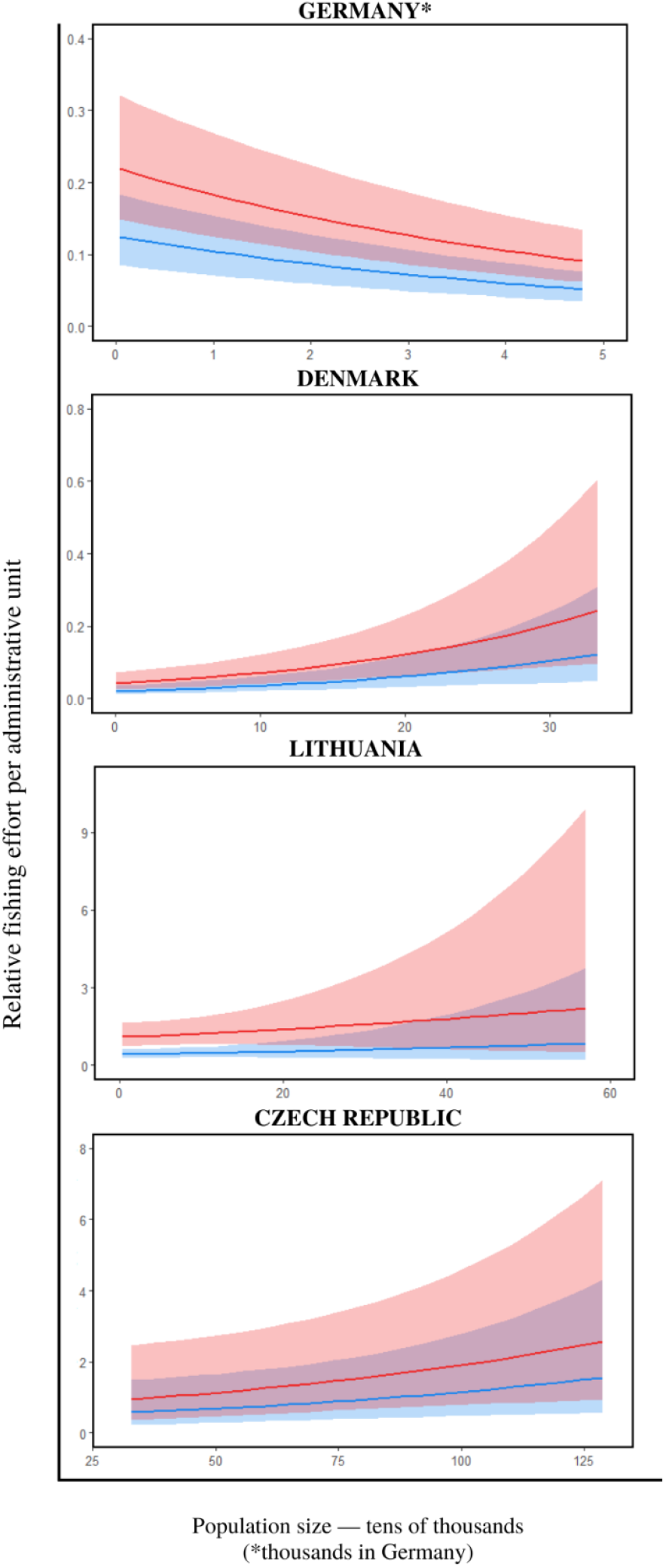
Association between relative fishing effort per administrative unit and human population density for pre lockdown study period. Associations remained similar for all five studied periods (see Fig. S8).

In summary, our study suggests an approximate doubling of recreational fishing effort across Europe in response to COVID-19 lockdown, with concomitant important societal, economic and ecological consequences. On one hand, recreational fishing has multiple human well-being and economic benefits, which have potentially provided valuable positive contributions to local economies and societies during the challenging years of the pandemic. On the other hand, such increases in effort are likely to represent a substantial increase in fishing pressure on inland and coastal ecosystems and species, many of which are already strongly affected by climate change and pollution (52–55). While increases in fishing effort of this magnitude in marine fisheries are not uncommon, this change typically occurs over a period of decades, rather than days and weeks (56). The implications of such acute changes in fishing effort are difficult to predict, but the short period over which these increases in effort occurred suggest that existing regulations may have been insufficient at maintaining fishing mortality within sustainable limits. In many cases, the inland and coastal populations that are targeted by anglers are unassessed (57, 58), so the potential impacts will remain unknown. However, closer collaboration between recreational anglers, smartphone application companies and scientists could greatly improve this situation, as digital technologies provide rapid and effective ways to collect data on fish population status.

## Material and Methods

### Data

Sonar data comprised individual daily sonar use events, identified through a unique encoded user ID, and including the time and geographical coordinates of the point at which the sonar device was first used on a given fishing trip. “Trips” often consisted of multiple readings at slightly different locations from the initial point of use. For each new reading, the user could select to either start a new trip or continue the same trip. We treated all trips by one user conducted within a single 24-hour period as one trip. We then used the starting coordinates of each trip to assign it to a single, country-relevant administrative unit: municipalities, in the case of Lithuania and Denmark, regions (Kraje) in the case of the Czech Republic, and districts (Landkreis) in Germany. This assignment allowed us to assess the relationship between angling effort and human population sizes. To ensure that only trips to officially recognised water bodies were included in the analysis (thereby excluding minor water bodies such as backyard dams and pools), we filtered the data for each country through a spatial layer of water bodies obtained through the GRPK – Spatial data set of (geo) reference base cadastre (open source data) from the Ministry of Agriculture of the Republic of Lithuania, the package RCzechia (59), for the Czech Republic, and the Esri Deutschland Open Data Portal (60) for Germany. We applied a buffer of 100 m around each water body for this data filtering to ensure that all shore anglers were included. Data from Denmark posed two challenges that required a different approach. First, there are over 120,000 lakes (> 0.01ha), many of which are not included in the Danish spatial layer of water bodies (61). A visual inspection of the data set revealed few (<10) instances of trips that were recorded away from water bodies (e.g., in buildings – presumably to test the device), so the data were not filtered by water bodies. Moreover, unlike in other countries, many device users in Denmark were sea anglers who fished offshore (25% of all recorded trips within Denmark). We included these anglers in the analysis by creating a 10 km buffer of spatial polygons along the coast, based on the closest land polygon, delimited by the spatial layer of municipalities in Denmark. This approach allowed us to assign each sea angling trip to the closest municipality of origin. All data processing was conducted in R (v4.2.1) (62), using *tidyverse* (v1.3.2) (63) for data manipulation and *sf* (v1.0.8) (64) for spatial analyses.

To obtain an indication of device users versus the total number of all anglers in a country (including those that fish only occasionally), we collated relevant information for each country. In Germany the total number of anglers in each administrative unit was estimated based on questionnaires published in Allensbach (65); in Denmark this estimate was based on an omnibus survey (66); in Czech Republic all anglers must register with an angling club and the numbers are available through a register maintained by the Ministry of Agriculture (67). In Lithuania anglers can purchase annual, monthly and one-day licences with, respectively, around 75 thousand, 20 thousand and 140 thousand sold annually during 2018-2021 (Ministry of Environment, Lithuania). It is not clear how many anglers purchase several monthly or one-day licences, although price differences were small (14 euros for annual, 5e for monthly and 1.4e for daily), so it is likely that monthly and daily licences are purchased by anglers who fish only occasionally. In addition, children under 16 and retirees do not require fishing licences, but fishing is a popular activity among retirees. Given that the assessments of angler numbers in other countries includes both regular and occasional anglers, we used an approximate estimate of 250 thousand anglers for Lithuania.

### Spatial and temporal COVID-19 restrictions

Government-imposed COVID-19 restrictions during the pandemic varied among and within European countries (Table 1). To ensure a consistent methodology, we used the Our World in Data compilation of ‘Policy Responses to the Coronavirus Pandemic’ curated by Ritchie et al. (68), available at: https://ourworldindata.org/policy-responses-covid (last accessed on 2022-10-17). This compilation of restrictions was collected and standardised by the Oxford Coronavirus Government Response Tracker (OxCGRT) (69) and has been used in other similar studies on human response to the COVID-19 pandemic (e.g. (9)). To define the lockdowns in our study, we used the daily index of ‘Stay-at-home requirements”, with any level of stay-at-home requirement (1-3) treated as a phase of ‘lockdown’. Based on this approach, each of the four countries had a pre-lockdown phase, starting on 2018-01-01, followed by two lockdown and two post-lockdown phases (Table 1, Fig. S1). The length of these phases differed slightly among countries, but the overall pattern was similar. We used a different approach for Denmark because Our World in Data compilation showed a protracted first lockdown (from March 20 to October 20, 2020), even though the Ministry of Health data in Denmark indicates none or very limited restrictions during the summer of 2020. For this reason, we followed the methodology of Gundelund and Skov (20) and defined the first lockdown from March 10 to May 31, 2020, and the second lockdown starting on October 23, 2020. The start of the second lockdown in Denmark was gradual and accelerated during autumn, but we choose October 23 as the starting point because this is when the government allowed up to 10 people to assemble (Danish Ministry of Health, 2022). The end of the second lockdown followed Our World in Data compilation, but we ignored a one-week (from 2020-11-16 to 2020-11-22) pause during Denmark’s second nationwide lockdown because it was likely too short to impact fishing effort. For all countries, the end of the second post - lockdown phase and the last day of our data was on 2021-12-31.

### Statistical analysis

We fit a Generalised Linear Mixed Model (GLMM) to data from each country with the phase of lockdown (pre lockdown, during lockdown 1, post lockdown 1, during lockdown 2, post lockdown 2), day of the week (weekday, weekend), administrative unit population size, the number of unique sonar users, and interactions between the phase of lockdown and day of the week as fixed effects. Random terms included season (spring, summer, autumn, winter), year (2018-2021) and national administrative region. The number of administrative units in each country was: Czech Republic 14, Denmark 96, Germany 374, and Lithuania 58 (Table S1). Lithuania and Denmark have more units, but some didn’t have any sonar device users recorded during the study and were removed from the analysis. Data were initially modelled with a Poisson-distributed response, which assumed that the variance was equal to the mean. Overdispersion occurs when the observed variance in the data is higher than the expected variance from the model, and we tested the dispersion ratio of models using the *performance* package (v0.9.2.4) (70). All Poisson models were overdispersed, so we instead assumed a negative binomial error structure.

The final model fitted to the data for each country took the form:

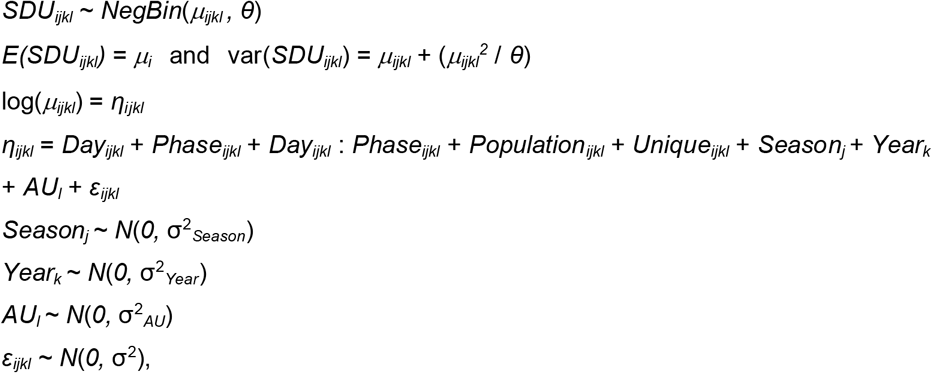

Where *SDU*_*ijkl*_ is the number of sonar device users on day *i* in *Season j* of *Year k* in administrative unit (*AU*) *l*, assuming a negative binomial distribution with mean *μ*_*ijkl*_ and variance *μ*_*ijkl*_ + (*μ*_*ijkl*_^2^ / *θ*). The parameter *θ* is the dispersion parameter. *Day* is a categorical variable for weekend/weekday and *Phase* for the phase of Covid-19 lockdown. *Population* is a continuous variable representing population density for the administrative unit. The variable *Unique* is a cumulative count of unique sonar device users on each day and was included in the model as a covariate to accommodate the increase in new sonar device users over time. The random intercept *Season*_*j*_ was included in the model to introduce a correlation structure between observations for different administrative units in the same season (spring: March to May, summer: June to August, autumn: September to November, winter: December to February), with variance σ^2^_*Season*_ distributed normally and equal to 0. Similarly, *Year*_*k*_ was included to accommodate correlation in the data between observations for different administrative units in the same season in the same year (2018-2021), and *AU*_*l*_ for correlation between observations for the same administrative units in the same season in the same year, while *ε*_*ijkl*_ is the residual variance in the model, with an assumption of normality and mean equal to 0.

We estimated the change in angling effort due to the lockdowns by comparing two estimates of the total effort in each phase of lockdown: an “observed” estimate that used model parameters that were derived from the phase in question, and a “null” estimate that assumed no lockdown by relying on model parameters (Table 2) from the pre lockdown phase. All details of these calculations are available together with the analysis codes in the GitHub repository.

## Data availability statement

The raw data were generated by users of the Deeper platform. Anonymous data was obtained through a data-sharing agreement. Requests for raw data should be directed to Vincentas Vienožinskis at Deeper. Derived data and codes for all the analysis are available on GitHub repository https://github.com/astaaudzi/covid_angling.

## Acknowledgments

This study has received funding from European Regional Development Fund (Project No. 01.2.2-LMT-K-718-02-0006) under grant agreement with the Research Council of Lithuania (LMTLT).

## Supplementary material

### Data fitting and exploration

Prior to model fitting, data exploration was undertaken following the protocol described in Ieno & Zuur (72). The data were examined for outliers in the response and explanatory variables, homogeneity and zero inflation in the response variable, collinearity among explanatory variables and the nature of relationships between the response and explanatory variables. The proportion of zeros in the number of sonar device users was high; Czech Republic 39%, Denmark 98%, Germany 87%, Lithuania 50%, though a high proportion of zeros does not necessarily indicate zero inflation in the data (Warton 2005). Zero-inflated models failed to improve model fit based on AIC, and a plot of the expected distribution of zeros against observed values using the *DHARMa* package showed close correspondence, further indicating a lack of zero inflation in the data. After fitting, the model validation protocol of Zuur & Ieno (72) was undertaken. Model assumptions were confirmed by plotting residuals against fitted values, covariates in the model, and covariates excluded from the model, which indicated no problems of model mis-fit.

**Table S1.** Summary of random effects for negative binomial GLMMs fitted to sonar buoy user data for the four countries investigated in the study; *σ*^2^ is the mean random effect variance for each model; τ_00_ is the model between-subject variance for the random effects year, season and administrative unit; ICC is the intra-class correlation coefficient, which is a measure of the degree of correlation within groups; N indicates the number of levels for the random effects year, season and administrative unit.

| Coefficient | Czech Republic | Denmark | Germany | Lithuania |
| --- | --- | --- | --- | --- |
| $\sigma^2$ | 0.81 | 3.68 | 2.93 | 1.06 |
| $\tau_{00}$ year | 0.27 | 0.17 | 0.04 | 0.06 |
| $\tau_{00}$ season | 0.31 | 0.08 | 0.14 | 0.04 |
| $\tau_{00}$ administrative unit | 0.49 | 1.07 | 2.22 | 0.77 |
| ICC | 0.57 | 0.26 | 0.45 | 0.45 |
| N<sub>year</sub> | 4 | 4 | 4 | 4 |
| N<sub>season</sub> | 4 | 4 | 4 | 4 |
| N<sub>administrative unit</sub> | 14 | 96 | 374 | 58 |

**Figure S1.**
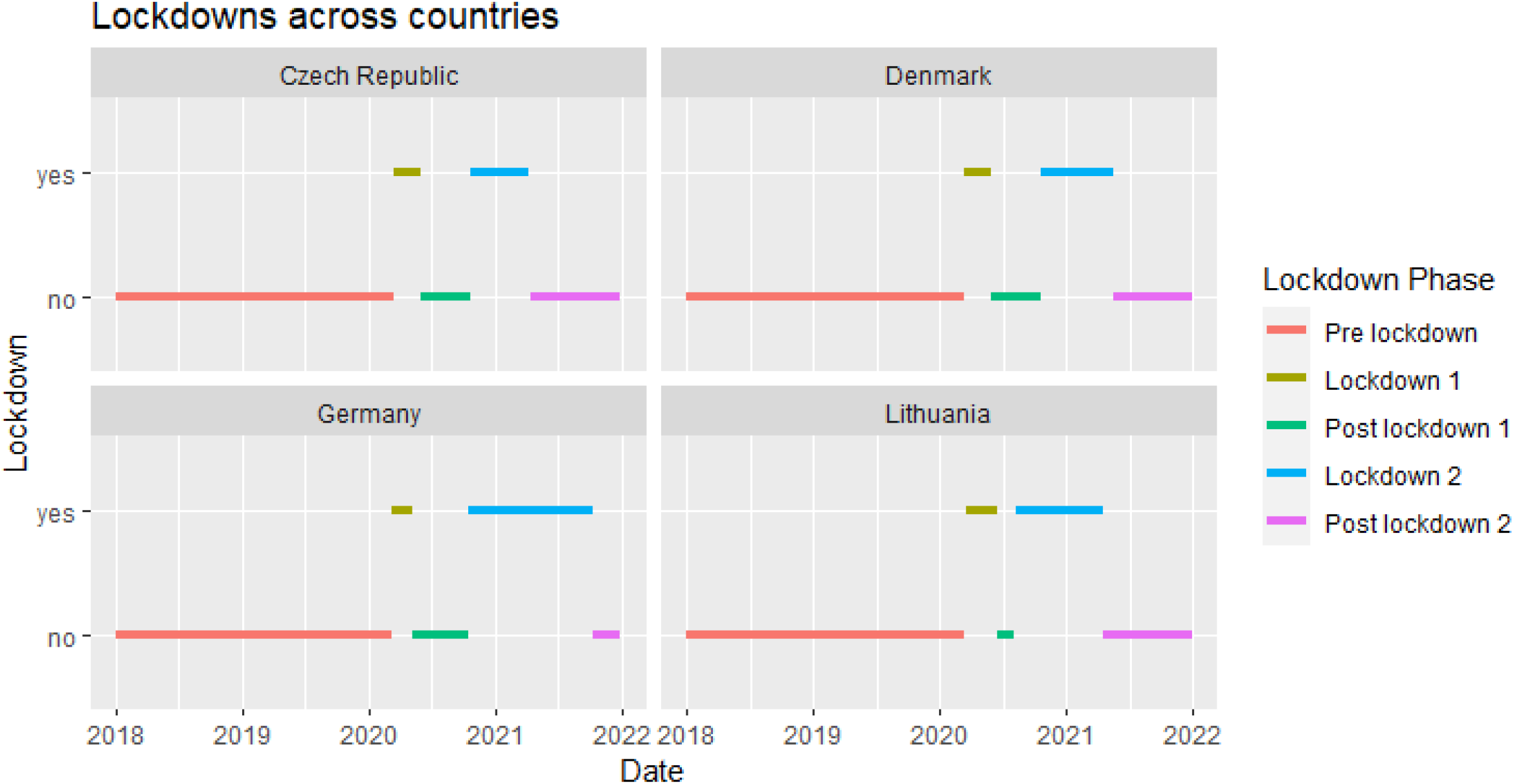
Dates and duration of lockdown phases, based on the daily index of “Stay at home requirements” by the Coronavirus Government Response Tracker, for the Czech Republic, Germany, and Lithuania; and on government data, according to the methodology described in Gundelund and Skov (20), for Denmark.

**Fig. S2.**
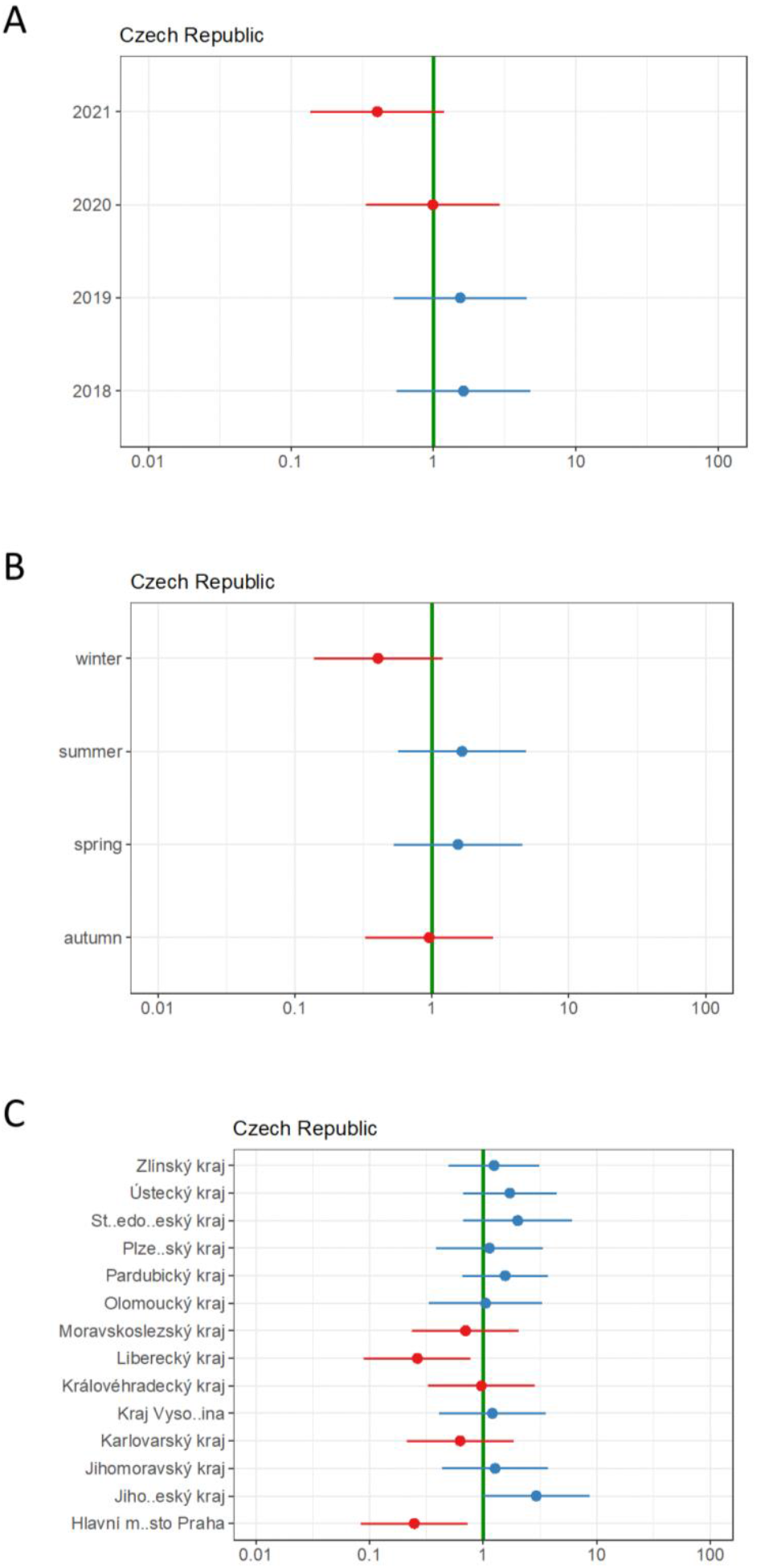
Random effects: Year, season, and administrative unit (*kraje*) in Czech Republic

**Fig. S3.**
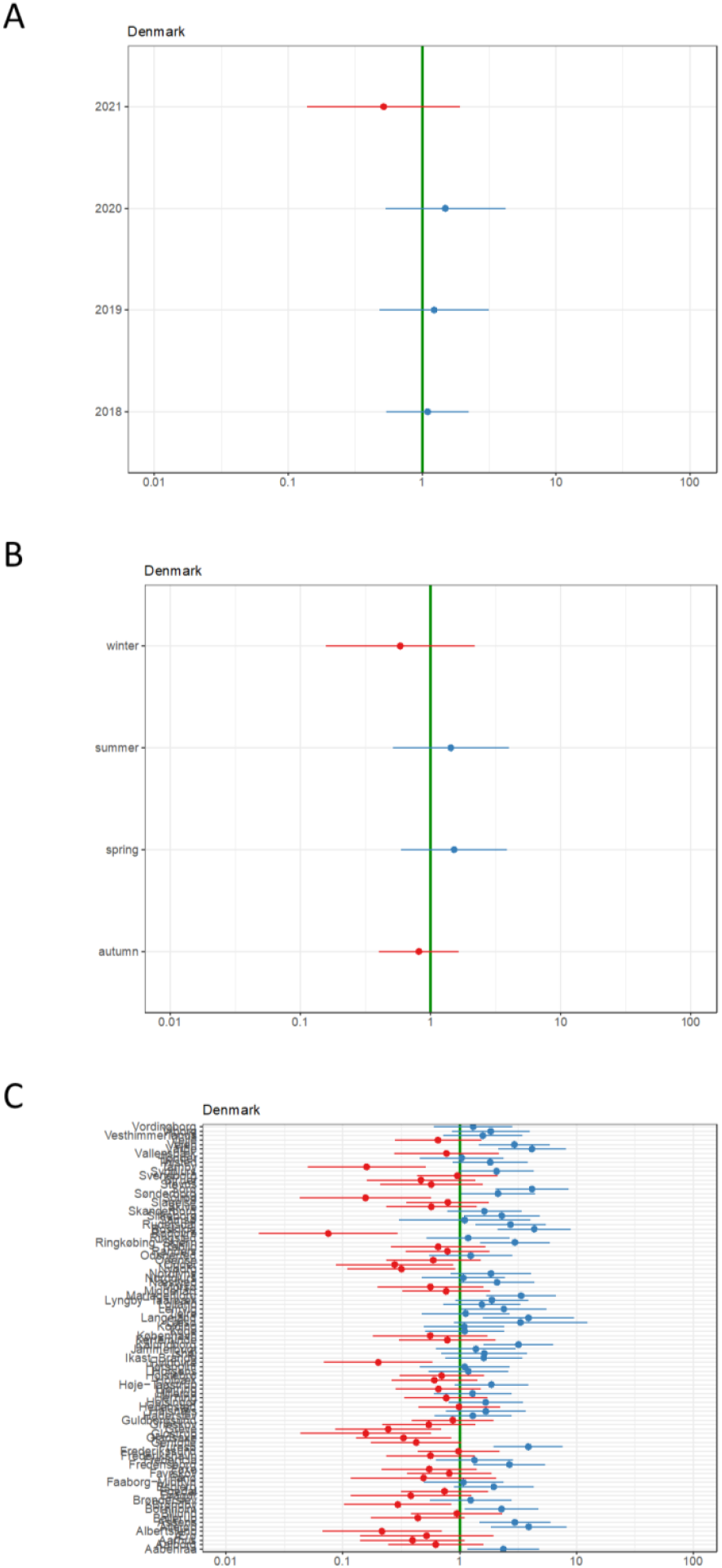
Random effects: Year, season, and administrative unit (*municipality*) in Denmark

**Fig. S4.**
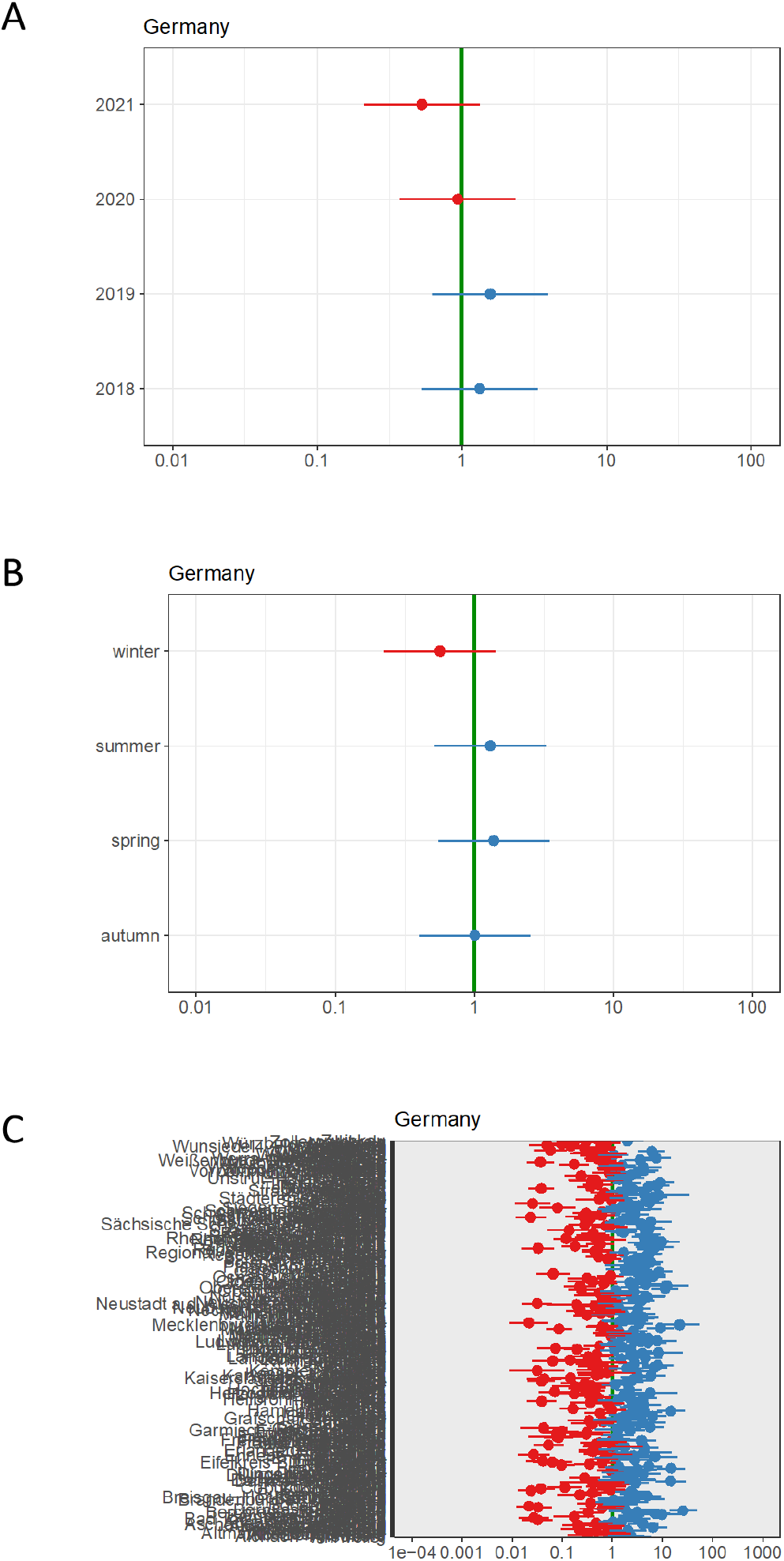
Random effects: Year, season and administrative unit (*Landkreis*) in Germany

**Fig. S5.**
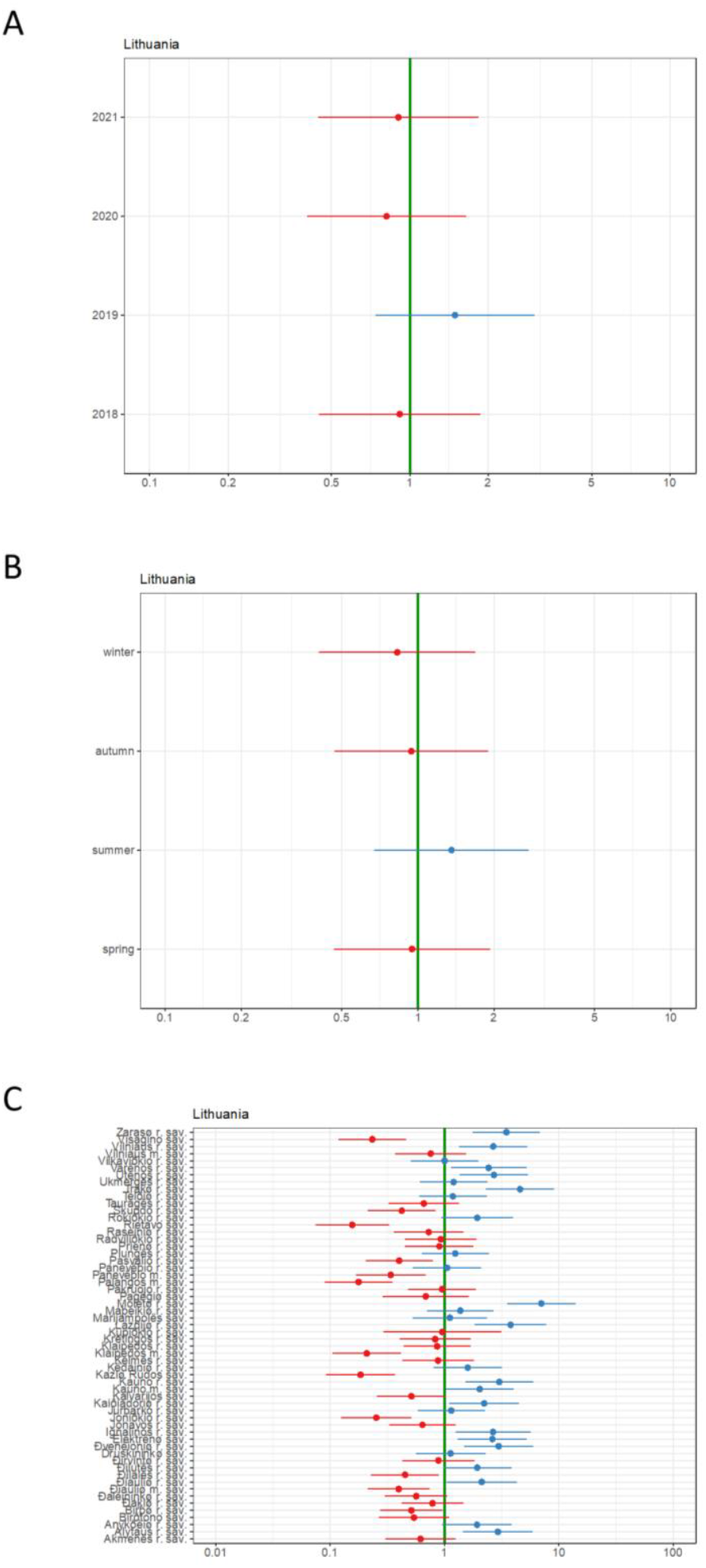
Random effects: Year, season, and administrative unit (*municipality*) in Lithuania.

**Fig. S6.**
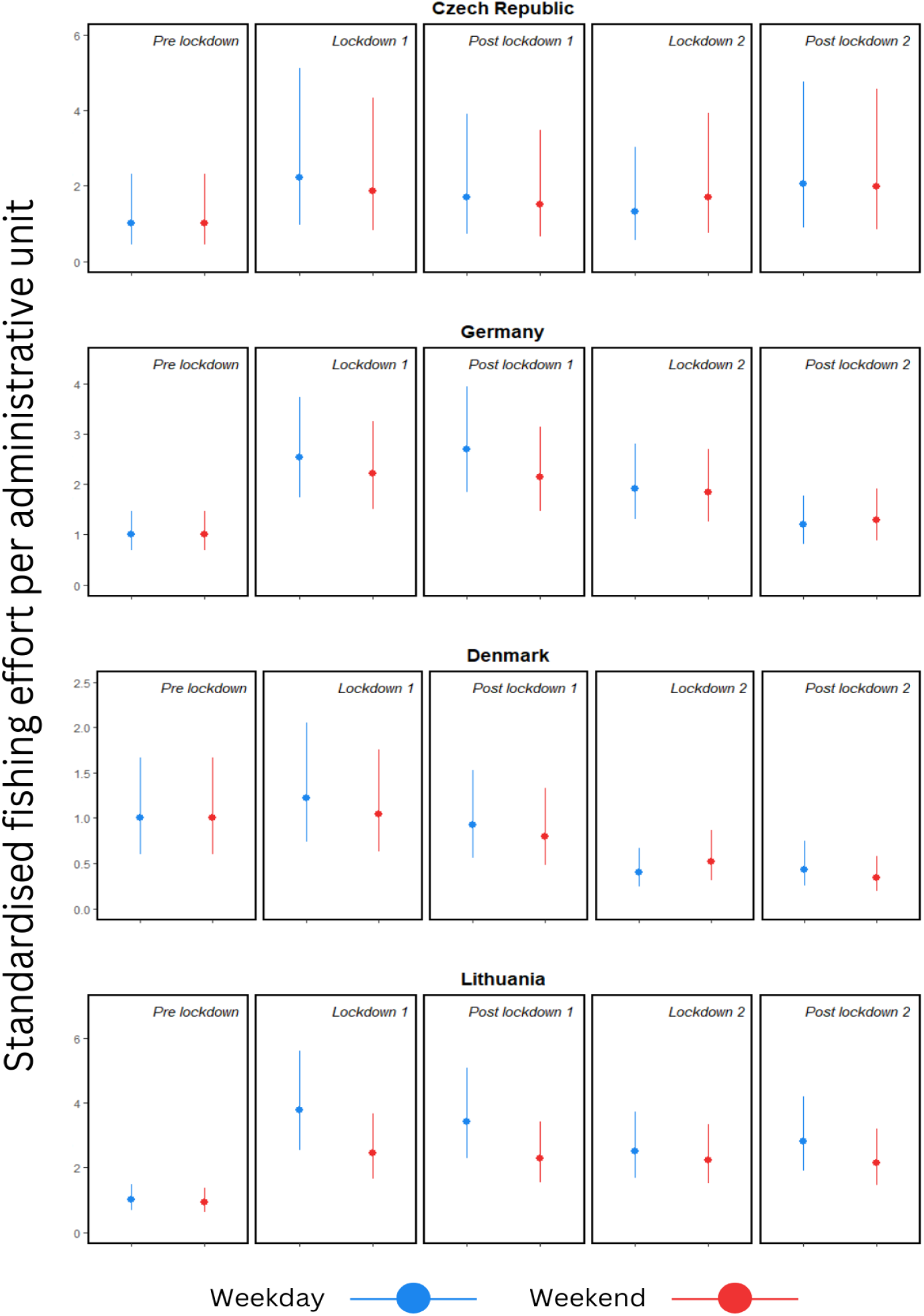
Estimated effort per administrative unit (as the number of sonar users during weekdays and weekends) for each of the five studied periods (Pre lockdown, During lockdown 1, Post lockdown 1, During lockdown 2, Post lockdown 2). The effort is estimated for an average administrative unit size for each country, scaled to the Pre lockdown stage for **both** weekdays and weekends.

**Fig. S7.**
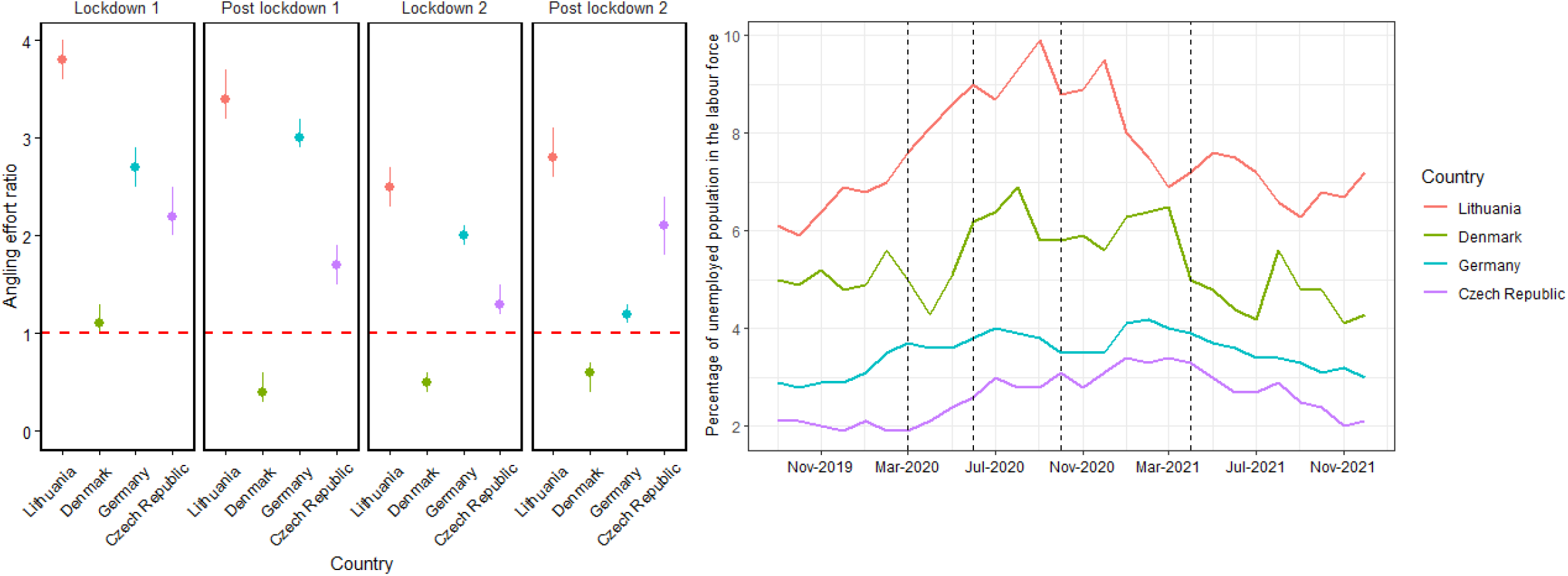
On the left, mean (+ 95% CI) estimated recreational fishing effort in each phase of lockdown in each of four countries as a ratio of the estimated fishing effort for each period assuming that lockdown had not been imposed. The horizontal red line at a ratio of 1 indicates no change in fishing effort in response to lockdown. Points above the line indicate increased fishing effort, points below the line indicate reduced fishing effort. On the right, variation of the percentage of unemployed population in the labour force across the same period. Approximate dates for the change of lockdown phases are indicated by dashed lines

**Fig. S8.**
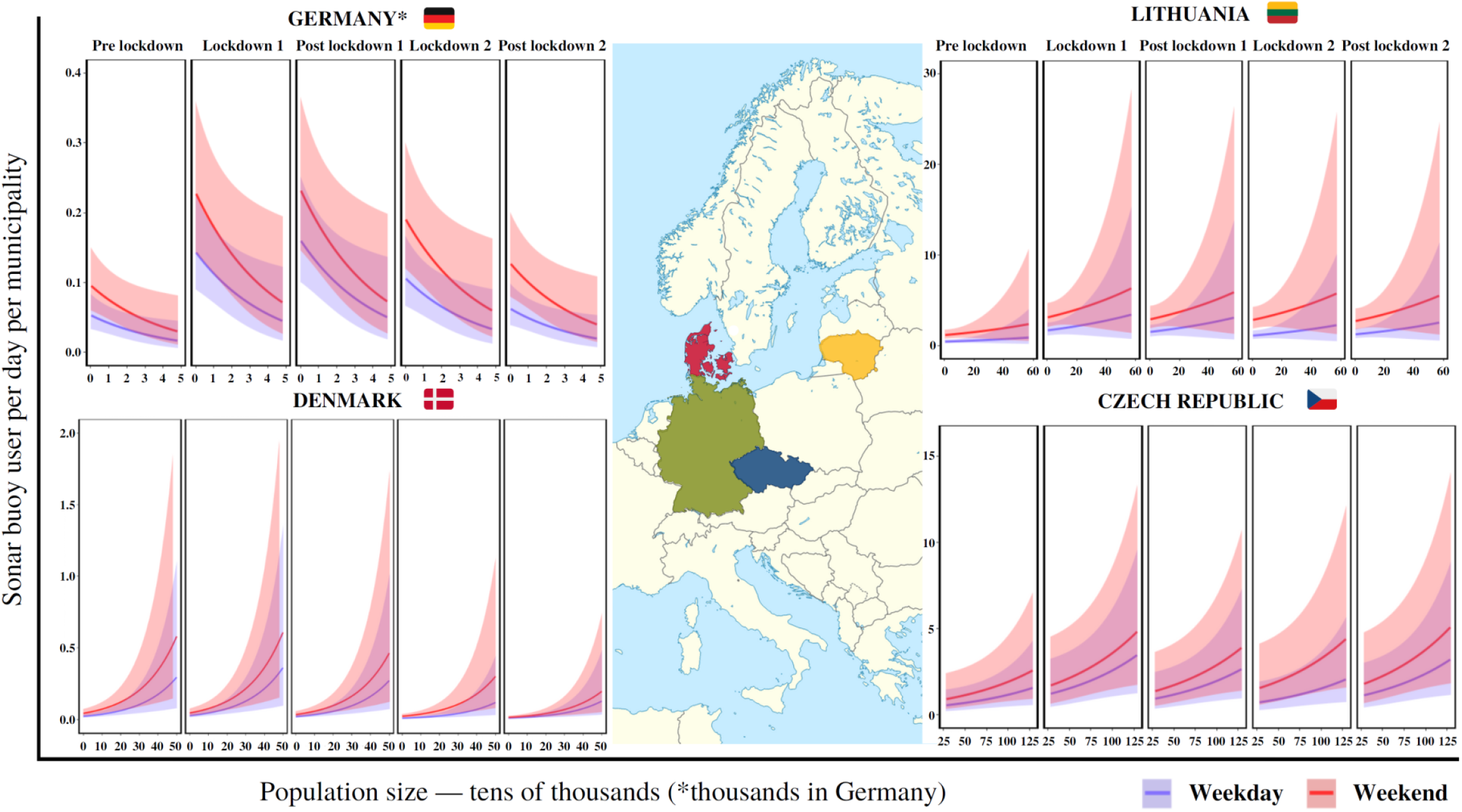
Estimated effort per administrative unit and population size by country for each study period (Pre lockdown, During lockdown 1, Post lockdown 1, During lockdown 2, Post lockdown 2). Effort is shown as the daily number of sonar users per administrative unit with different population sizes.

